# An estimation of the absolute number of axons indicates that human cortical areas are sparsely connected

**DOI:** 10.1101/2021.06.07.447453

**Authors:** Burke Q. Rosen, Eric Halgren

## Abstract

The tracts between cortical areas are conceived as playing a central role in cortical information processing, but their actual numbers have never been determined in humans. Here we estimate the absolute number of axons linking cortical areas from a whole-cortex diffusion-MRI (dMRI) connectome, calibrated using the histologically-measured callosal fiber density. Median connectivity is estimated as ~6200 axons between cortical areas within-hemisphere and ~1300 axons inter-hemispherically, with axons connecting functionally-related areas surprisingly sparse. For example, we estimate that <5% of the axons in the trunk of the arcuate and superior longitudinal fasciculi connect Wernicke’s and Broca’s areas. These results suggest that detailed information is transmitted between cortical areas either via linkage of the dense local connections or via rare, extraordinarily privileged long-range connections.

## Introduction

The major tracts connecting cortical areas have long been central to models of information-processing in the human brain [1]. Such models have been refined and applied to development and disease with the advent of diffusion-MRI or dMRI [2]. However, dMRI only provides *relative* connectivity, not the absolute number of axons. Relative connectivity is very useful in many circumstances but more constraints would be possible if the absolute number of axons connecting different cortical areas could be estimated. Here we describe and apply a novel method for translating from dMRI-derived streamlines to axon counts.

The ratio between these two measures is obtained by comparing the number of streamlines pass through the corpus callosum to the number of axons, as measured with histology. The corpus callosum is uniquely well-suited for this purpose. Electron microscopy (EM) can be used to unambiguously count the total number of callosal axons because the limits of the callosum are well-defined, axons are aligned, and sections can be cut perpendicular to their axis. Likewise, dMRI-derived streamlines can be unambiguously and exhaustively assigned to the callosum because the source and destination of all axons passing through the callosum as a whole are well-defined. In contrast to the ipsilateral fiber tracts, in which the majority of axons enter or leave the tract at point between the fascicular terminals, commissural axons must all be connecting the two hemispheres. Fortunately, despite these differences, the mean and variability of cross-sectional axon density of the corpus callosum is quite similar to that of telencephalic white matter in general [3–5], permitting the streamline:axon ratio calculated from callosal fiber to be applied to intra-hemispheric connections.

It has long been recognized that as the number of cortical neurons increases, maintaining the same probability of connectivity between neurons would require that axon number increase approximately with the square of neuron number, and this would require too much volume, impose an unsustainable metabolic load [6], and actually decrease computational power due to conduction delays [7]. The consequent imperative to minimize long distance cortico-cortical fibers has been posited to be reflected in exponential decline in cortical connectivity with distance [8], and to be partially compensated for with a small-world graph architecture [9], granting special properties to rare long-distance fibers in a log-normal neural physiology and anatomy [10]. However, this organizing principle is rarely explicitly addressed in terms of individual axon counts, nor in a manner both granular and exhaustive with respect to cortical areas. In our histologically calibrated dMRI-derived estimation of this intercortical axon counts, we find that the widespread cortical integration implied by behavioral and mental coherence, and routinely observed in widespread physiological synchronization, belies a surprising small absolute number of long-range axons connecting cortical areas.

## Materials and Methods

The basic principle of our method is very simple. Given a dMRI-based measure of total inter-hemispheric connectivity in arbitrary units and the physical number of axons traversing the corpus callosum, the conversion factor between the two can be obtained by dividing the first by the second. Specifically, we started with the total inter-hemispheric tractography strength reported in our dMRI connectome of the Human Connectome Project (HCP) cohort [11]. For each individual, the cross-sectional area of the corpus callosum was obtained using the standard FreeSurfer structural MRI pipeline [12]. Multiplying this number by a histologically ascertained callosal fiber density [3] yields an estimate of the number of axons traversing an individual’s corpus callosum. Dividing this count by individuals’ inter-hemispheric dMRI streamline value yields the conversion ratio from the arbitrarily-scaled dMRI metric to the absolute number of axons. Note that this procedure is independent from the scale of the dMRI metric, requiring only that it be proportional to the absolute number of fibers. A moderate proportionality has been observed in comparisons of dMRI and retrograde tracing in macaque [13,14]. Therefore, while the ratio itself is study-specific, the procedure can be applied to any dMRI tractography method or parameter set, provided that the dMRI method returns a continuous distribution of connectivity values and has reasonably similar sensitivity to callosal and ipsilateral fiber tracts. To demonstrate this, we repeated the analysis using data from an alternate tractography of the HCP cohort [15]. For detailed methodology, see S1 Appendix: Extended Methods and Results.

## Results

Our previous dMRI study [11] includes estimated tractography streamlines between all 360 parcels of the HCP-MMP1.0 atlas [16] for each of the 1,065 individuals in the HCP dMRI cohort. The sum connectivity between the 180 left hemisphere cortical parcels and the 180 right (180^2^ parcel pairs) constitutes the total callosal connectivity, on average 1.25×10^8^ streamlines. Based on our assumed fiber density of 3.7×10^5^ axons/mm^2^ [3] and measured callosal cross-sectional area (mean = 689.45mm^2^), we estimate this cohort to have 2.6×10^8^ callosal fibers on average. The mean quotient between these two quantities is 2.00 axons-per-streamline, a ratio specific to the dMRI methodology and parameters used.

Applying the conversion factor to the inter-parcel connectivity from our prior study yields an estimate of the absolute number of axons connecting different cortical areas (Fig 1A), 2.43×10^9^ axons in total. This implies that <22% of the ~11.5×10^9^ cortical pyramidal cells [17,18] project outside their parcel with the remainder being short-range horizontal or U-fibers. Furthermore, because 51% of inter-areal axons are to adjacent areas, <11% of pyramidal cells project beyond the adjacent parcel. Axons are approximately log-normally distributed among non-adjacent parcel pairs, with adjacent pairs having disproportionally high axon counts (Fig 1B). Median connectivity is ~6,200 axons between cortical areas in the same hemisphere, and ~1,300 inter-hemispherically. The number of axons in the median *inter-*hemispheric connection is ~20% of that in the median *intra-*hemispheric connection, similar to that found with histological tracing in macaques [19]. The number of callosal axons is not significantly affected by subject sex, though subjects ages 26-30 have slightly more inter-hemispheric axons than subjects ages 22-15 (S3 Fig). This sparse long-range connectivity is consistent with its exponential fall-off measured with dMRI in humans and histologically in other mammals [8,11] and with previous statistical estimates based on traditional neuroanatomy [20]. In comparison to the number of axons that would be necessary for complete interconnectivity of neurons in different cortical areas with each other, the estimated number is ~10^11^ times smaller (see S1 Appendix for calculation).

**Fig 1.**
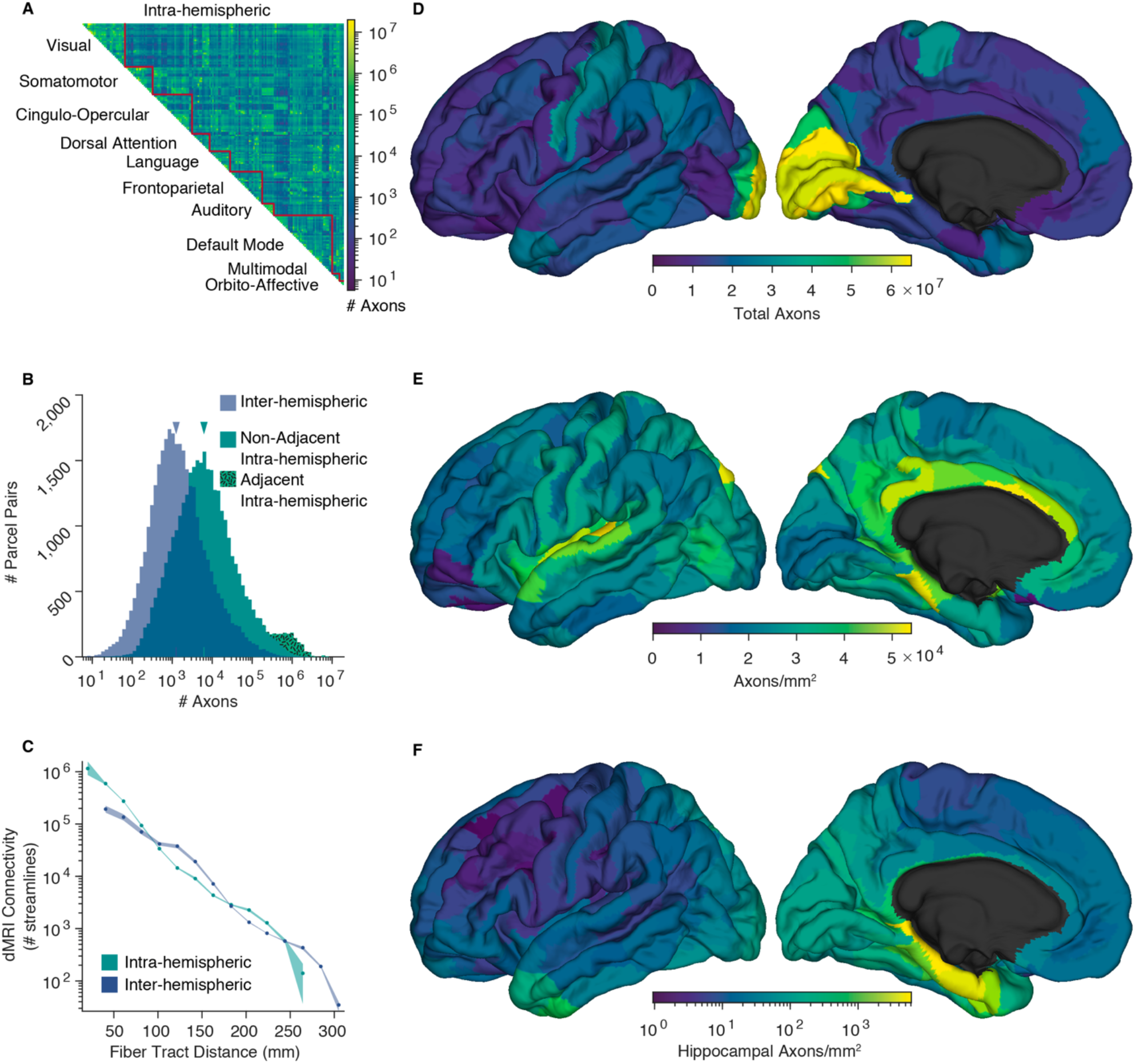
The number of axons estimated to interconnect the 360 cortical parcels of the HCP-MMP1.0 atlas. **(A)** Connectivity matrix of intra-hemispheric axon counts, averaged across the two hemispheres. Parcels are ordered into 10 functional networks. **(B)** Histograms showing the distribution of inter- and intra-hemispheric pairwise axon counts. Physically adjacent and non-adjacent parcel pair intra-hemispheric histograms are stacked. Median connectivity, indicated, is ~6,200 axons between cortical areas in the same hemisphere, and ~1,300 inter-hemispherically. **(C)** Comparison of intra- and inter-hemispheric dMRI connectivity as a function fiber tract distance. Pairwise values averaged within 15 fiber-length bins. Shading shows bootstrapped 95% confidence intervals. **(D-F)** axon counts and densities averaged across the two hemispheres and visualized on the left fsaverage template cortex [12]. **(D)** total axons connecting each parcel to all others. **(E)** axons connecting each parcel to all others, normalized by the reference parcel’s area. **(F)** axons connecting the hippocampus to the rest of the cortex, normalized by the area of the cortical parcel, shown in log scale. The source data for this figure can be found at https://doi.org/10.5281/zenodo.6097026.

Our method requires that dMRI-tractography performs in a roughly similar manner when applied to intra-versus inter-hemispheric connections. We evaluated this similarity by comparing linear regressions of distance-matched log-transformed streamlines, and found no difference between the slope or intercept of inter- and intra-hemispheric connections (Fig 1C); within the common distance domain a paired r-test showed no difference in correlation, *p* = 0.58. The total count of a parcel’s axons to or from all other areas is much less variable when normalized by the parcel’s area (Fig 1D,E). Multiplying the total number of fibers by the effective fiber cross-sectional area (the inverse callosal packing density, 2.7×10^−6^ mm^2^/axon) yields 6.6×10^3^ mm^2^ of cortex or 3.7% of the total white-gray interface. Note that the effective axonal area includes myelin, supporting cells and intercellular space in addition to the axon proper. The cross-sectional packing density of human prefrontal white matter is quite similar to callosal values with an average of 3.5×10^5^ myelinated axons/mm^2^ [4] after correction for tissue shrinkage, and this varies among prefrontal regions by a less than factor of 2 [5]. The percent of fiber-allocated cortical area is quite similar to the ~4% of total cortical fibers Schüz and Braitenberg estimated [20] are contained in the corpus callosum and long fascicles. The remaining ~95% of the cortical gray-white interface area is likely occupied by the short range U-fiber system which is difficult to assess with dMRI.

We estimated the fraction of axons traversing the entire length of the arcuate / superior lateral fasciculus (AF/SLF) between termination fields centered on Broca’s and Wernicke’s areas. The total number of AF/SLF axons was derived by multiplying the tracts’ cross-sectional areas by mean axon density of ipsilateral tracts. Using published estimates of the cross-sectional areas of 160.6 mm^2^ and 51.5 mm^2^ for the left and right AF and 213.8 mm^2^ and 174.4 for the left and right SLF [21], and a shrinkage-corrected axon density of 3.5×10^5^ axons/mm^2^ [4], yields a total of 1.3×10^8^ and 0.8×10^8^ axons in the left and right AF/SLF. These values were compared to the number of tractography-derived axons connecting AF/SLF termination fields [16] according to consensus definitions from reference [22], see S4 Fig. When the anterior termination field, centered on Broca’s area is defined as HCP-MMP1.0 parcels 44, 45, 6r, IFSa, IFSp, and FOP4 and the posterior termination field, centered on Wernicke’s area as parcels PSL, RI, STV, and PFcm, trans-terminal axons account for only 0.6% and 0.8% of tract axons in the left and right hemispheres. If the termination fields are liberally expanded to also include parcels 47l and p47r in Broca’s and parcels PF, PFm, and PGi in Wernicke’s then these percentages increase to a still modest 1.9% and 2.9% of total tract axons. Please note that in this calculation all axons between Broca’s and Wernicke’s areas are assumed to pass through the AF/SLF; to the degree that some pass outside the AF/SLF, our estimates should be decreased.

The volume of hemispheric white matter (WM) occupied by axons between cortical parcels is equal to the sum of all axons’ lengths multiplied by their cross-sectional areas. The mean fiber tract lengths of connections were taken from our prior dMRI analysis [11]. For the effective cross-sectional area of axons we again assumed an effective cross-sectional area equivalent to the inverse of the callosal fiber density [3]. So calculated, the volume occupied by cortico-cortical and hippocampo-cortical fibers is 4.3×10^5^ mm^3^, or about 96% of the total MRI-assessed WM volume. This implies that the number of long-range fibers cannot be larger than our estimate unless the axon density calculated from the corpus callosum histology is mistakenly low, but this is a linear effect that would need to be unrealistically inaccurate for it to change our main conclusions. Another possibility is that axon density was correctly measured for the corpus callosum but is higher for intra-hemispheric fibers *and* the number of axons per streamline is higher intra- than inter-hemispherically. These both need to be the case because there is no space available in the hemispheric white matter to contain more axons unless they are smaller. However, the histological data indicates that axonal density is approximately the same intra- and inter-hemispherically [3,4] and the comparison of distance-matched streamlines (Fig. 1*C*). suggests that the number of axons per streamline intra- and inter-hemispherically is also approximately equal. The remaining WM volume may seem insufficient for connections with subcortical structures. However, the major subcortical structures known to communicate with the cortex (thalamus, amygdala, striatum, nucleus basalis, locus coeruleus, etc.) contain, in sum, less than 1% of the number of cortical neurons [23–25], see S1 Appendix: Extended Methods and Results. Even under the unlikely assumption that all excitatory thalamic neurons project to the cortex with a density across all cortical parcels proportionally to their area, their axons would only comprise ~4% of the total WM volume.

We conducted a series of simulations examining the consequences of possible errors in axonal packing density and axon-to-streamline ratio. As noted above, these parameters are co-constrained by the physical volume available for white matter. At our derived ratio and assumed packing density, the volume of inter-areal axons is just under the observed total white matter volume. Large net errors in the assumed packing density would result in total inter-areal axon volumes that are inconsistent with the hemispheric white matter volume, a value that is well established. Consequently, in our simulations, axonal packing density and axon-to-streamline ratio were reciprocally changed in order to maintain a constant total cerebral white matter volume. Since our parameter estimates are based primarily on fibers passing through the callosum, which are longer in general than intra-hemispheric fibers, parameter inaccuracies are more likely for short fibers. In order to evaluate the effect of such inaccuracies, we first simulated the effect of assuming that dMRI sensitivity is systematically reduced (and thus axon count underestimated) for shorter fibers. Uniformly doubling the number of axons with lengths < 40 mm in this way yields only a 37.5% increase in the total number of inter-areal axons, and because rank-order of observations is not altered the median axon counts are unchanged.

Conceivably, our parameter estimates are accurate for the longest fibers (which tend to pass through the callosum) but are progressively less accurate for shorter inter-areal distances. Consequently, we simulated assuming that our packing density and dMRI sensitivity estimates at the longest fiber lengths were as in our base model, and then increased the dMRI streamline-to-axon ratio linearly as the fibers got shorter. Axon density was adjusted to maintain white matter volume constant. Even with unrealistically large systematic and matched errors in the two estimates, median inter-areal axon count are only modestly increased, see S5 Fig. For example, adjustments resulting in twice the axon density and half the sensitivity of dMRI to axons at short inter-parcel distances increase the median number of inter-parcel axons by only ~36%. As reviewed below, errors in inter-areal axon count due to miss-estimation of axonal packing density are likely to be relatively unbiased with respect to inter-areal fiber length, and reported measures vary less than a factor of two. Our simulations indicate that such errors would have only marginal effects on the median number of inter-areal axons.

In order to demonstrate that the key principles and findings of this report are robust to the details of tractography procedure, we repeated our analysis on the tractography data of Arnatkeviciute and colleagues [15]. These data consist of a 972 subject subset of the HCP cohort and use the same parcellation but a different method to reconstruct the fiber streamlines. These data contain fewer streamlines and the estimated numbers of axons per connection are broadly comparable, being somewhat fewer but within an order of magnitude of our primary estimate. We find 8.4×10^8^ total axons and 1.1×10^8^ callosal axons per subject on average, with medians of ~1100 and ~130 axons for pairwise intra- and inter-hemispheric connections between cortical areas (S2 Fig), i.e., even more sparse than those calculated from our tractography data. These data confirm that, independent of tractography procedure, cortical areas are sparsely connected.

## Discussion

In this study we estimated the absolute number of axons interconnecting cortical areas by calibrating dMRI-based tractography using the histologically ascertained cross-sectional fiber density of the corpus callosum and found that long-range cortico-cortical connections are quite sparse. Our method depends on histological estimates of callosal axon packing density and leverages the unique properties of the callosum. It assumes proportionality between the number of axons connecting two areas and the number of dMRI tractography streamlines for the areal pair, and it assumes approximate parity between dMRI sensitivity to inter- and intra-hemispheric connections. In order to estimate the volume occupied by inter-parcel cortico-cortical connections, we make the further assumption that the effective axonal packing density is reasonably uniform across the hemispheric white matter.

Although there are few reports of axonal density in the corpus callosum [3,26], they are consistent with each other and with reports of axonal density of intra-hemispheric tracts [4]. Our study relies on the data reported by Aboitiz et al. [3], because it is the most systematic count of which we are aware. There is a later study, [26], which provides a value only slightly lower, 2.83×10^5^ vs 3.7×10^5^ axons/mm^2^ after correction for tissue shrinkage [27]. However, since this later study was primarily a survey of axon diameter with packing density only incidentally recorded, it is conceivable that density values derived from could be small underestimates. Mis-estimated tissue shrinkage is possible but likely to be <10% and callosal areas were measured with a well-validated and widely-used in vivo method [12]. The axon count estimate does not require that the packing densities of the corpus callosum and ipsilateral white matter be the same, but rather the lesser assumption that the axon to dMRI streamline ratio be reasonably uniform across the various instances of long distance cortico-cortical connectivity. However, if the packing densities of ipsilateral and callosal long distance connections are also similar then the total hemispheric white matter volume provides an absolute constraint on the number of long distance connections, and this provides a powerful validation of our estimate.

The literature provides converging evidence that the fiber packing density of white matter varies by at most a factor of 2 across the cortex, including the corpus callosum. A histological study of human cortex found only a 7% difference in the axon densities of the callosum versus the superior longitudinal and inferior occipitofrontal fasciculi [26]. Zikopolous and Barbas [4] found that the cross-sectional packing density of human prefrontal white matter is remarkably similar to callosal values with an average of 3.5×10^5^ myelinated axons/mm^2^ after correction for tissue shrinkage and that this varies among prefrontal regions by a less than factor of 2 [5]. In addition, while dMRI-based estimates of axon density and caliber are imperfect, they suggest that axon density varies by less than 2-fold both between major ipsilateral tracts and within each tract along their length [28]. While packing density is not directly equivalent to axon caliber, the two are likely inversely related. Axon diameter, as estimated with dMRI, varies by 20% among ipsilateral tracts and by at most a factor of 2.2 among white matter voxels, including those of the corpus callosum [29,30]. Overall, distributions of dMRI-derived axon diameter for the callosum [31] and whole cerebrum [30] are similar with the bulk of values between 2.5 and 5 μm. Histological measurements in macaques concur that axon diameters are very similar within the callosal and non-callosal segments of major fasciculi and vary by less than 2-fold across the cortex [32]. Furthermore, even this limited variation in diameter is not systematically dependent on the length of axons but rather on their regions of origin and termination [33].

Concerning dMRI-to-axon-count proportionality, it has been shown that there is a moderate linear correlation between dMRI-traced streamlines and the number of fibers identified with histological tracers connecting cortical regions in macaques [13,14] and a strong correlation in ferrets [34]. Based on these data, Donahue and colleagues [13] concluded that dMRI tractography was capable of quantitively describing cortico-cortical white matter tracts with approximately order-of-magnitude precision using a high-quality dataset such as the HCP. In line with previous statistical estimates of whole-cortex axon counts based on traditional neuroanatomy [20], the numbers of axons in pairwise connections are probably correct within an order of magnitude. Estimates of this precision are useful as we find that that inter-areal connectivity derived from dMRI varies over more than seven orders of magnitude.

While we derive a single dMRI-to-axon count factor, it is likely that the true conversion ratio varies somewhat among connections due to local microstructural differences other than axon count such as axon caliber, packing density, or myelination. However, the major fasciculi (including the corpus callosum) have similar axon calibers and packing densities, varying by only a factor of ~2 across the cortex when examined in humans histologically [4,5,26] or with dMRI [28–30] and histologically in macaques [32]. More generally, packing density is a function of local cellular interactions, especially with oligodendrocytes, and because these are the same in the corpus callosum and intra-hemispheric white matter, the *a priori* expectation is that packing density would be similar, as histological data supports. Myelination has only a modulatory effect on dMRI-detected anisotropy, with most of the effect derived from axonal membranes [35]. Nevertheless, microstructural variation in dMRI-to-axon count ratio may be a source of noise in our estimates. Our simulations of miss-estimation of dMRI-to-axon ratio and packing density show that these errors only marginally affect estimated axon counts and do not alter our conclusions. Our findings were also similar when we repeated our analysis on an alternative, somewhat sparser dMRI tractography dataset [15], demonstrating that the details of the tractography algorithm do not affect the overall tenor of our results. While the scope of this study is limited to long-range fibers, we note that shorter, more superficial U-fibers are systematically less myelinated and of lesser caliber than their inter-areal counterparts [5] and therefore the procedure outlined here may require modification before being applied to them.

As previously stated, this methodology assumes a reasonable degree of parity in the sensitivity of the dMRI tractography to intra- and inter-hemispheric fiber tracts. Consistent with this assumption, we found little difference between distance-matched dMRI connectivity for callosal versus ipsilateral connections. If callosal axons were more easily detectable, this would be reflected as an upward displacement of the inter-hemispheric trace (blue) above the intra-hemispheric trace (green) across the entire distance domain in Fig 1C, which is not evident. This parity is perhaps unsurprising because over most of their trajectories, inter-hemispheric fibers are subjected to the same crossing fiber issues as intra-hemispheric. Specifically, the corpus callosum is a distinct tract for only about 15-35 mm, but its fibers range in length up to about 300 mm. Thus, the fraction of an inter-hemispheric tract that resides within the callosum is inversely proportional to its total length. Consequently, if there were enhanced detection of callosal fibers as streamlines by dMRI, one would expect the inter-hemispheric (blue) trace in Fig 1C to be elevated primarily at short fiber lengths and depressed at long fiber lengths, resulting in a noticeably steeper slope for the inter-hemispheric than the intra-hemispheric trace, which is not observed. These data support the applicability of the scaling factor derived from inter-hemispheric fibers to intra-hemispheric fibers.

The corpus callosum was used to calibrate the estimate because of its unique properties: it has a well-defined cross-sectional area, more than ~99% of inter-hemispheric cortico-cortical axons are routed through it [3,36], and essentially no fibers leave or enter the tract between the two hemispheres. This is in sharp contrast to non-commissural fasciculi. While it may be commonly assumed that within major cortical fasciculi the majority of axons terminate or originate at the ends of the tract and thus carry information along its entire length, an alternative conception is that these large bundles are composed mostly of axons shorter than the total fascicular length which enter and exit the tract at various points. By analogy, the former assumption likens a tract to a tunnel, where all traffic is trans-terminal, whereas the latter conceives of tracts as like interstate highways, where very few vehicles travel the entire route. In a supplementary analysis we compared these models for the arcuate / superior lateral fascicular (AF/SLF) system. The total number of AF/SLF axons was estimated using the tract diameters [21] and packing densities [4] taken from the literature. The number of trans-terminal axons was determined by defining termination field parcels centered on restricted and inclusive definitions of Broca’s and Wernicke’s areas [22]. Depending on the assumptions, only about 1-5% of the axons in a middle section of these fasciculi are trans-terminal. These percentages may be over-estimates since they assume that all fibers between the posterior and anterior areas travel through the AF/SLF. However, even if these values are a 2-fold underestimate, it suggests that only a small fraction of the axons in the ipsilateral fasciculi are trans-terminal. The evidence indicates that the ‘highway’ model is more apt for the ipsilateral fiber tracts and this conception is consistent with neural wiring being driven, in large part, by exponential distance rules [8]. This of course, does not apply to the corpus callosum, as there is no inter-terminal cortex to project into.

Importantly, the reconceptualization of major intra-hemispheric tracts as containing few fibers connecting their distant terminals is still consistent with the long- and well-established impression from blunt dissection [37] and dMRI orientation maps [38] that the hemispheric white matter is largely composed of well-defined long-distance tracts. Indeed, we estimate that ~96% of the white matter is composed of inter-areal axons. What these observations suggest is that the major tracts arise from the tendency of axons to grow in alignment with axons which are already present using established mechanisms of adhesion and fasciculation [39]. Thus axons are free to join tracts at various points, and tend to proceed together (fasciculate), but again are free to leave whenever they approach their own target. In other words, axons are joined in a given tract because they share a direction rather than an origin and destination.

The inter-areal axon counts we derive here permit other interesting quantitative estimates which may inform models of cortical neurophysiology. For example, the connections between Wernicke’s and Broca’s areas are thought to integrate receptive and expressive aspects of language, but we estimate that there are only ~58,000 axons between the core cortical parcels in these regions (parcels 44 and PSL), fewer than two for each word in an average university student’s vocabulary [40]. Another example where quantitative appreciation of direct axonal connections may influence neuro-cognitive models are the hippocampo-cortical interactions subserving recent memory, which are commonly posited to carry information regarding the contents of the memory trace during memory formation, consolidation and retrieval. We estimated that areas distant from the hippocampus, notably the dorsolateral prefrontal cortex, may be connected to it by <10 axons/mm^2^ (Fig 1F) including both efferent and afferent axons, yet hippocampo-prefrontal interactions are considered crucial for contextual recall [41]. We estimated the average neural density in the cortex as ~92,300 neurons/mm^2^ by dividing the 16.34×10^9^ cortical neurons (including interneurons) from [17] by the 1.77×10^5^ mm^2^ mean white—gray surface area of the HCP cohort used. If the sparse connectivity suggested by our calculations is correct, it implies that hippocampo-dorsolateral prefrontal interactions in memory are likely mediated by polysynaptic pathways.

These constraints encourage consideration of models of cortical function where connections are dense but mainly local, i.e. a small world network with intense interconnections within modules and sparse projections between them. While this general principle is widely accepted, the scale of the vast gulf in absolute connectivity between local and long-range connections is startling. This network architecture provides for the wide and efficient distribution of information created by local processing within modules [8,9]. A more uniformly connected cortex would require more white matter, necessitating a more voluminous cerebrum and the human cortex is near the limit after which an increase in size reduces computational power [6,7]. Functionally, a deep reservoir of weak connections enables a large number of states and eases state transitions [10]. Modeling suggests that long range covariance and even synchrony can be achieved through activation of multi-synaptic pathways rather than direct connections [42,43], and possible signs of these have been observed experimentally in humans [44,45].

While the long-range direct cortico-cortical axons are few in number, we note that axons are heterogeneous and that these counts are a limited proxy for true inter-areal connectivity. Axons, especially those connecting architectonically similar regions, may have a disproportionate impact on the flow of information despite their rarity [46–48], by virtue of their morphology (e.g. greater diameter, larger termination fields, greater axonal arborization, or more numerous *en passant* varicosities) or by molecular synaptic specializations. For example, while only ~5% of the synapses to V1 layer 4 come from the lateral geniculate body in the macaque [49], they have an outsized effect on their firing [50]. The importance of rare inter-module connections might also be enhanced if they are focused on a small location within cortical parcels (i.e. the rich club [51]), but this has not been convincingly demonstrated. Lastly, it is useful to note that these quantitative considerations are radically different in other species, where the smaller number of cortical neurons and shorter inter-areal distances allow greater connectivity between cortical areas, as well as a larger proportion of subcortical connectivity [7].

## Acknowledgements

This work was supported by National Institutes of Health grants RF1MH117155 and R01NS109553 (EH). Data were provided, in part, by the Human Connectome Project, WU-Minn Consortium (Principal Investigators: David Van Essen and Kamil Ugurbil; 1U54MH091657) funded by the 16 National Institutes of Health (NIH) Institutes and Centers that support the NIH Blueprint for Neuroscience Research; and by the McDonnell Center for Systems Neuroscience at Washington University. The funders had no role in study design, data collection and analysis, decision to publish, or preparation of the manuscript.

## Data availability

The processed data used in this study are available at https://doi.org/10.5281/zenodo.6097026. Datafiles are in Matlab v7.3 format. The Human Connectome Project’s raw imaging data and FreeSurfer outputs may be downloaded from https://db.humanconnectome.org.

## Code availability

The main standalone analysis Matlab code used in this study is available at https://doi.org/10.5281/zenodo.6097026. Additional non-standalone supporting code will be made available upon request.

## S1 Appendix: Extended Methods and Results

### Diffusion MRI

The diffusion-weighted MRI (dMRI) connectivity data used to estimate inter-parcel axon totals was described in our prior report [1] whose data are available in full at https://doi.org/10.5281/zenodo.4060485. Preprocessed imaging data for 1,065 healthy young adults were retrieved from the Human Connectome Project WU-Minn consortium 1200 release [2] (https://db.humanconnectome.org). We segmented the cortex into 180 parcels per hemisphere following the HCP-MMP1.0 atlas [3] and performed probabilistic tractography using the probtrackX tool [4] from FSL [5] using tractography parameters detailed in our prior report [1]. This analysis yielded a 360 × 360 dMRI connectivity matrix for each subject which provided the example data for applying the method presented here. For probtrackX-derived connectivity, the raw unit of dMRI connectivity strength is the number of streamlines connecting seed and target regions. The absolute magnitude of this metric is arbitrary as it scales with the number of samples drawn from the probability distribution of diffusion parameters in each voxel, in our analysis 5,000.

The individual pairwise streamline counts and fiber tract length matrices for the replication analysis were described in [6] whose data are available at https://doi.org/10.5281/zenodo.4733297. These data consist of a 972 subject subset of the 1,065 WU-Minn HCP subjects included in our primary analysis. Arnatkeviciute and colleagues similarly segmented the cortex following the HCP-MMP1.0 atlas [3]. In contrast to our primary dataset, tractography was performed with MRtrix3 [7] using second-order integration over fiber orientation distributions (iFOD2) [8], see [6] for details. When confined to the same subject subset and non-null connections, the pairwise tract lengths and log-transformed streamline counts are highly correlated (r^2^ = 0.78 and r^2^ = 0.69, respectively) between Arnatkeviciute et al. [6] and our study [1]. The rate of exponential decay in the number of streamlines vs fiber tract distance is also similar, λ = 26.1 mm in their study vs 23.4 mm in ours, though less connectivity variance is explained by distance in their results (r^2^ = 0.12 vs. r^2^ = 0.52). Results for these tractography data are shown in S2 Fig.

### Cross-sectional area of the corpus callosum, white matter volume, and parcel areas

The HCP WU-Minn consortium 1200 release [2] includes the standard outputs from the FreeSurfer `recon-all` pipeline [9]. With default parameters, this pipeline labels the 1 mm^3^ voxels of the `aseg.mgz` volume which contain the corpus callosum with a fixed 5mm lateral extent. Dividing the number of callosum-label voxels by 5 yields an estimate of its cross-sectional area, in mm^2^. The total white matter volume is reported in the `aseg.stats` output of `mri_segstats`. Parcel areas and adjacency were determined [10] on triangular mesh of each subject’s reconstruct white matter surface, or white matter – gray matter interface (`?h.white`). These data were used for both the primary and replication analyses.

### Cross-sectional fiber packing density in corpus callosum and other hemispheric white matter

Our estimate of callosal fiber density was based on the electron microscopy (EM) study by Aboitiz and colleagues [11] who reported a fiber density of 38 per 100 μm^2^ or 3.7×10^5^ axons/mm^2^ after accounting for tissue shrinkage to 96% volume after Epon fixation [12]. This is consistent with their finding of 2.65×10^5^ axons/mm^2^ as measured with light-microscopy in twenty brains as well, assuming shrinkage to 43% volume for paraffin fixation [12] and an estimated ~20% of fibers not detectable with light microscopy [11], as well as the 2.83×10^5^ non-exhaustively counted axons per mm^2^ reported in a more recent EM study of callosal axon diameter in two brains [13]. This value was derived by dividing the reported mean count of axons per callosal region by the callosal area surveyed, and correcting for EM preparation tissue shrinkage. An additional light microscopy study of callosal axon density in eleven control cortices reports an average of 1.12×10^5^ myelinated axons per mm^2^ [14], assuming shrinkage to shrinkage to 43% volume. This somewhat lower figure may be explained by the inclusion of four children’s cortices in their cohort, or by mis-assumed shrinkage, but all reports are within half an order-of-magnitude of each other.

### Distance matched intra- vs. inter-hemispheric dMRI connectivity

One assumption of the method described in this report is that the dMRI tractography methodology has approximately similar sensitivity to callosal and ipsilateral connections. This is difficult to determine without ground truth connectivity, as intra- and inter-hemispheric connections may have true anatomical differences. Nevertheless, we examined distance-matched intra- vs inter-hemispheric connections (Fig 1*C*) and found no statistical evidence that they are different. Pairwise connections were averaged in 15 distance bins, log-transformed and linear fits obtained for intra- and inter-hemispheric connections. For the overlapping distance domain, the intra-hemispheric slope is −0.30, 95% confidence interval = [−0.33 −0.26] with an intercept of 6.18 [5.91 6.45] and the inter-hemispheric slope is −0.26 [−0.29 −0.24] and intercept of 5.93 [5.72 6.14]; the 95% confidence intervals of both parameters are overlapping. In addition, a paired r-test of correlation differences [15] failed to reject the null hypothesis, *p* = 0.58. The parity between distance-matched intra- and inter-hemispheric connections also holds when the linear fit is obtained for over the entire distance domain, with an intra-hemispheric slope and intercept −.31 [−0.33 −0.28] and 6.24 [6.01 6.46], respectively; −.28 [−0.30 −0.25] and 6.00 [5.78 6.23] for inter-hemispheric connections. In the replication analysis with alternative tractography data [6] (S2 Fig C) the paired r-test likewise failed to reject the null hypothesis, *p* = 0.34. The intra-hemispheric slope and intercept are −.37 [−0.37 −0.78] and 4.19 [3.25 5.13], respectively; −.25 [−0.30 −0.20] and 3.30 [2.82 3.79] for inter-hemispheric connections.

### Estimating axon counts and axon volume from dMRI

From the histological estimate of callosal fiber density, MRI-derived measures of callosal cross-sectional area and white matter connectivity, the conversion factor between dMRI connectivity and physical axons can be estimated as follows:

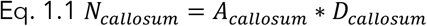

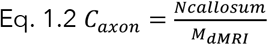

Where *A*_*callosum*_ is the cross-sectional area (mm^2^), *D*_*callosum*_ is the assumed fiber density (3.7×10^5^ axons/mm^2^) of the corpus callosum, *N*_*callosum*_ is the estimated total number of callosal fibers, *M*_*dMRI*_ is the magnitude of dMRI callosal connectivity, in our case, the number of tractography streamlines, and *C*_*axon*_ is the conversion factor from dMRI to physical fibers (axons/streamline).

In addition to connectivity strength, or number of streamlines, the dMRI tractography analysis also yields a mean streamline length for every parcel-pair connection. After converting dMRI connectivity strength to physical axon count as described above, the total volume of the estimated axons can be calculated as follows:

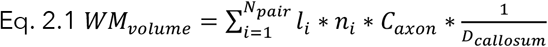

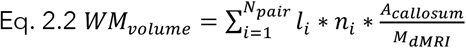

Where *WM*_*volume*_ is the estimated axonal white matter volume (mm^3^) and *N*_*pair*_ is the total number of parcel pairs. *l*_*i*_ and *n*_*i*_ are the length (mm) and number of dMRI streamlines in the *i*^*th*^ parcel pair. Eq. 2.1 can be simplified into Eq. 2.2 by substituting Eqs. 1.1 and 1.2 into 2.1.

### Parcel grouping into functional networks

The parcel ordering and the ten functional networks shown in Fig 1*A* were defined in our previous report [1] and were modified from twelve networks identified by Ji and colleagues in a resting-state fMRI study on the HCP cohort [16] by merging the primary and secondary visual networks as well as the ventral and posterior multimodal networks.

### Effects of sex and age on inter-hemispheric connectivity

In their histological study of 20 cortices, Aboitiz and colleagues [11] found no significant effects of sex or age on the total number of fibers, fiber density, and cross-sectional area of the corpus callosum. As we assume a constant fiber density, our estimate of the total number of inter-hemispheric fibers is a linear function of the callosal cross-sectional area. As the ages of the HCP 1200 cohort are grouped into four broad ranges in the open access data, we investigated the effects of sex and age on the estimated number inter-hemispheric axons with 2×4 fixed effects ANOVA, treating the age group factor as categorical. Mean and individual values are shown in S3 Fig. Note that while the number of pairwise axons is approximately log-normally distributed across areal pairs, it is approximately normally distributed across individuals.

We found that, on average, males have 2.61×10^8^ inter-hemispheric axons, 95% confidence interval = [2.58×10^8^ 2.64×10^8^] and females have 2.50×10^8^ [2.47×10^8^ 2.52×10^8^] axons. Subjects ages 22-25, 26-30, 31-35, and 36+ year old have 2.50×10^8^ [2.46×10^8^ 2.55×10^8^], 2.58×10^8^ [2.55×10^8^ 2.61×10^8^], 2.55×10^8^[2.51×10^8^ 2.59×10^8^], and 2.35×10^8^ [2.15×10^8^ 2.54×10^8^] axons, respectively. Intervals show bootstrapped 95% confidence. We find no effect of sex on the number of inter-hemispheric axons, *F*_1,1064_ = 3.471, *p* = 0.0627. Removing the one female outlier does not bring the mean difference into statistical significance, *F*_1,1063_ = 3.653, *p* = 0.0562. The interaction of sex and age on the number of inter-hemispheric axons is likewise not significant *F*_3,1064_ = 2.282, *p* = 0.0776. Age does significantly affect the number of inter-hemispheric neurons, *F*_3,1064_ = 5.100, *p* = 0.0017. Post-hoc pairwise tests show that only the difference between the 22-25 and 26-30 year old age groups is significant, *F*_1,1064_ = 7.646, *p* = 0.0058. These results are broadly consistent with those of Aboitiz et al. [11]. A more in-depth examination of the effects of age on corpus callosum requires a cohort with which includes more older subjects as well as more precise age values.

### Thalamo-cortical fiber volume estimation

The scope of our previous dMRI tractography study [1] was confined to cortico-cortical (including cortico-hippocampal) connectivity. In order to estimate the total volume of thalamo-cortical axons, we first calculated the number of excitatory thalamic neurons by summing the total number of neurons in the thalamus [17], excluding the reticular nucleus, zona incerta, limitans/suprageniculate, and subthalamus, and including from the remainder the ~62% proportion of thalamic neurons that are excitatory [18]. The number of such neurons is 22.6×10^6^. We assumed that each such thalamic neuron projects a single axon to the cortex, and we allocated these axons to cortical parcels in proportion to their area. The inverse callosal packing density was used for effective cross-sectional area, as with cortico-cortical connections. To estimate the length of thalamo-cortical fiber tracts, for each parcel *i*, we first identified the thalamic voxel nearest to parcel’s centroid. We then used the inter-parcel fiber tract length [1] from the centroid of parcel *i* to the identified parathalamic parcel centroid. To this figure we added the Euclidean distance from the parathalamic centroid to the nearest voxel of the thalamus, ensuring that estimated thalamo-cortical fiber tract length is always non-zero. As with cortico-cortical connections, the total white matter volume of each connection was estimated by taking the product of the number of axons, the effective cross-sectional area, and the estimated fiber tract length (Eq. 2).

### Estimation of proportion of actual inter-areal axons to the number that would be needed for complete inter-connectivity

We estimated that the total number of inter-areal axons is ~2.43×10^9^. Assuming n = 16.34×10^9^ total cortical neurons (including interneurons) [19], the total number of connections needed for complete whole-cortex connectivity is n(n-1) = 2.67×10^20^. We estimated the neural density of the cortex as ~92,300 neurons/mm^2^ by dividing 16.34×10^9^ by 1.77×10^5^ mm^2^ mean white—gray surface area of the HCP cohort used. The total number of connections needed for complete inter-connectivity within each area is approximately 1.18×10^18^, given by

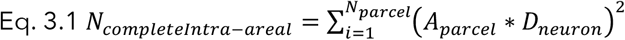

Where A_parcel_ is the area of each parcel and D_neuron_ the neural density. Subtracting the connections needed for within-area inter-connectivity from whole cortex interconnectivity yields 2.66×10^20^, which is 1.10×10^11^ times greater than the number of inter-areal contacts we calculate with our method.

### Number of neurons in non-cortical regions communicating with the cortex

The number of neurons in different human brain regions has been reviewed by Blinkov and Glexer [20] and von Bartheld et al. [21]. The number of neurons in the amygdala have been estimated as ~13×10^6^ [22], striatum as ~55×10^6^ [23], and thalamus as ~23×10^6^ (thalamocortical, see above). These are the only recipients or targets of cortical connections with significant numbers of neurons. Other locations, such those which are the origins of serotonin [24], norepinephrine [25], dopamine [26], or acetylcholine [27] afferents to the cortex all have ~50,000 axons each. Summed together, the number of cells in all of the above structures comes to ~0.5% of the number of cortical cells. Direct projections outside of the brain are exceedingly rare, with the number of Betz cells in the primary motor cortex [28] or medullary pyramid [29] both estimated as ~100,000. It is difficult to escape the conclusion that, at least in humans, the cortex communicates mainly with itself.

---

**S2 Fig:**
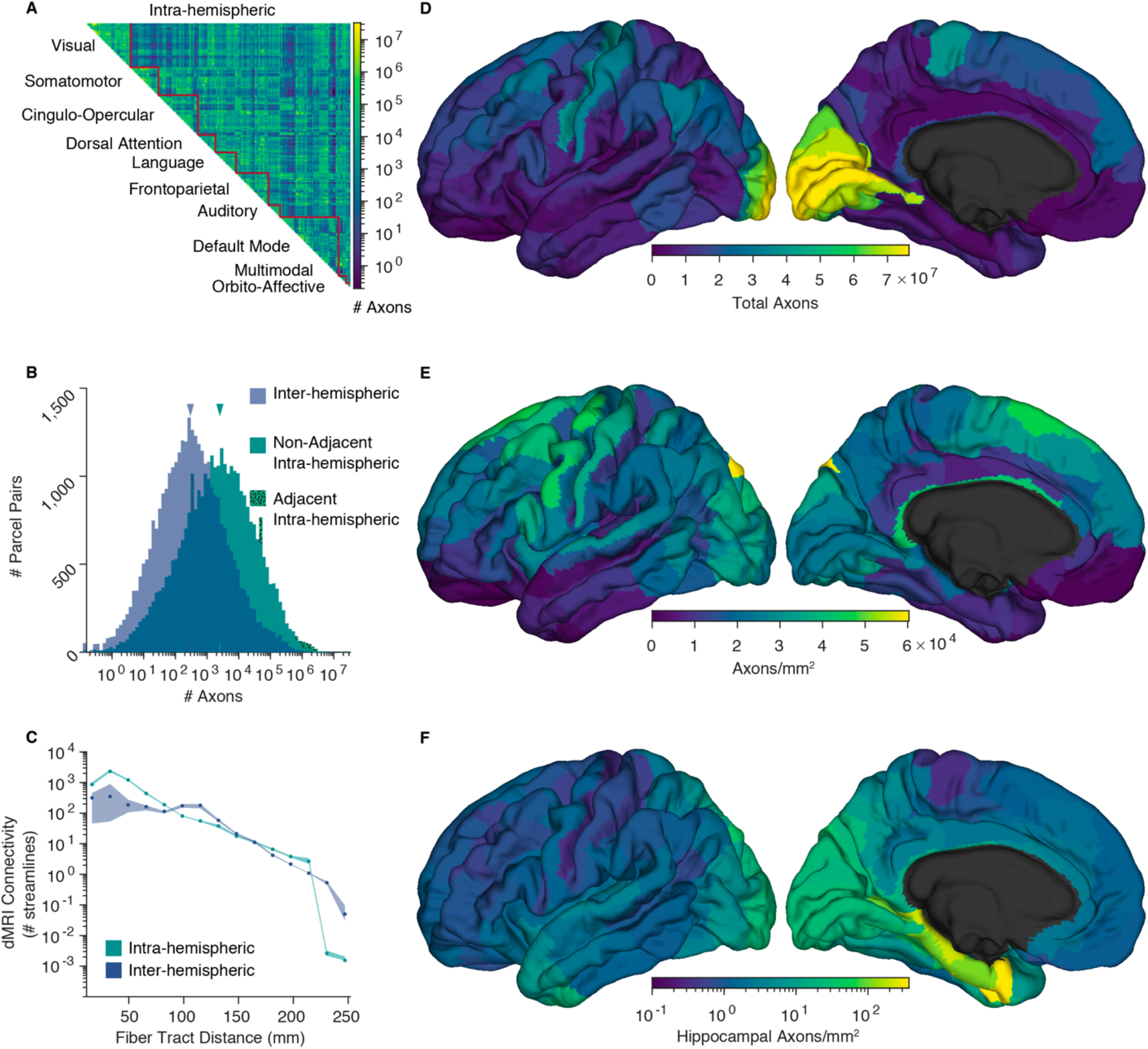
Replication results. Results obtained by repeating the analysis using an alternative tractography dataset [1] (A) Connectivity matrix of intra-hemispheric axon counts, averaged across the two hemispheres. Parcels are ordered into 10 functional networks. (B) Histograms showing the distribution of inter- and intra-hemispheric pairwise axon counts. Physically adjacent and non-adjacent parcel pair intra-hemispheric histograms are stacked. Median connectivity, shown in gray, is ~2,500 axons between cortical areas in the same hemisphere, and ~300 inter-hemispherically. Parcel pairs with zero axons connecting them are represented by the bars left of the y-axis. (C) dMRI connectivity as a function fiber tract distance. Pairwise values averaged within 15 fiber-length bins. Shading shows bootstrapped 95% confidence intervals. (D-F) axons counts and densities averaged across the two hemispheres and visualized on the left fsaverage template cortex [2]. (D) total axons connecting each parcel to all others. (E) axons connecting each parcel to all others, normalized by the reference parcel’s area. (F) axons connecting the hippocampus to the rest of the cortex, normalized by the area of the cortical parcel, shown in log scale. The source data for this figure can be found at https://doi.org/10.5281/zenodo.6097026.

**S3 Fig:**
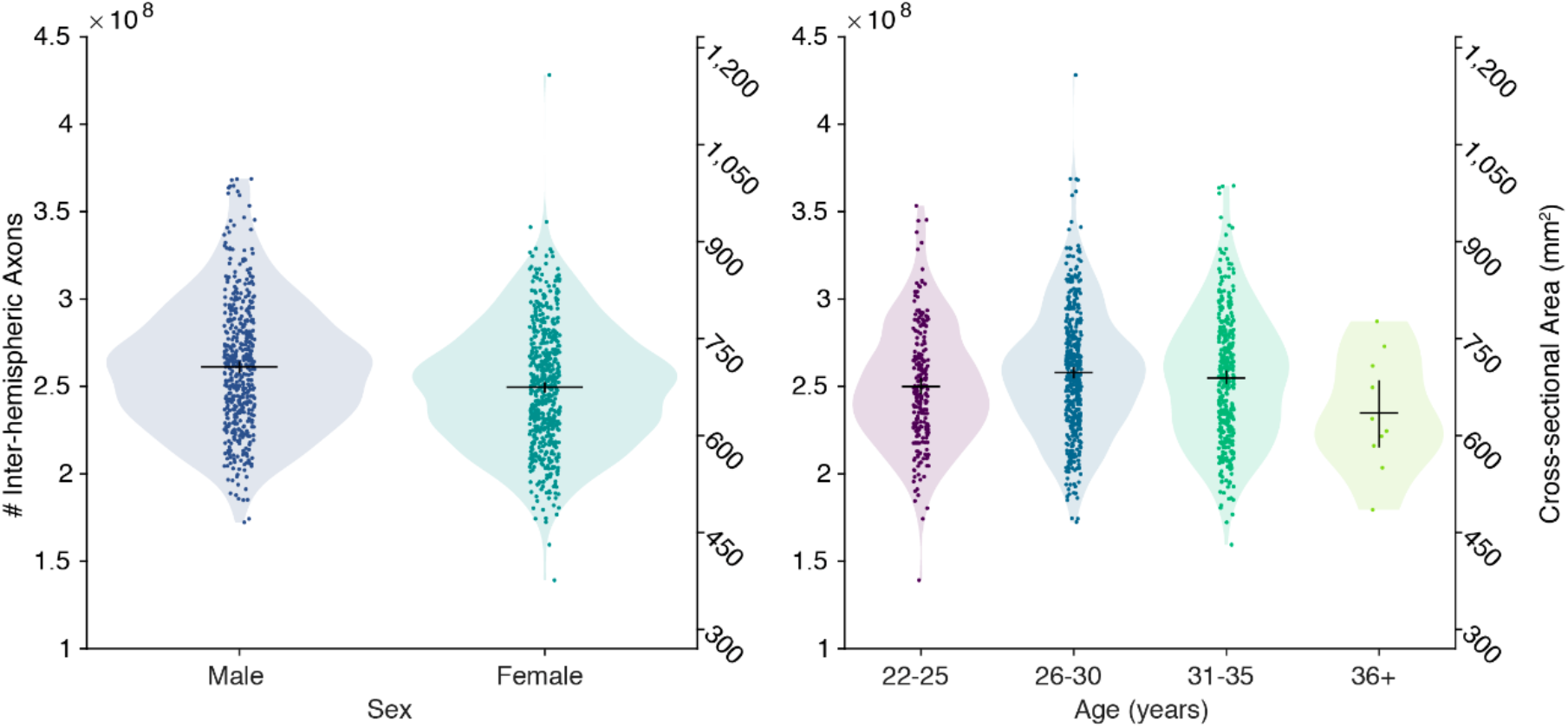
Effects of sex and age on the inter-hemispheric connectivity. Each individual’s total number of estimated inter-hemispheric axons is shown with a marker. The black horizonal bars show the group means and vertical bars the bootstrapped 95% confidence intervals these means. Shading shows a kernel density estimate of the group distributions. Note that while the number of pairwise axons is approximately log-normally distributed across areal pairs, it is approximately normally distributed across individuals. The only significant group difference is between the 22-15 and 25-30 age groups, *F*_1,1064_ = 7.646, *p* = 0.0058. Interactions between sex and age effect were not significant. As we assume a constant fiber density, our estimate the total number of inter-hemispheric fibers is a linear multiple of the callosal cross-sectional area. The source data for this figure can be found at https://doi.org/10.5281/zenodo.6097026.

**S4 Fig:**
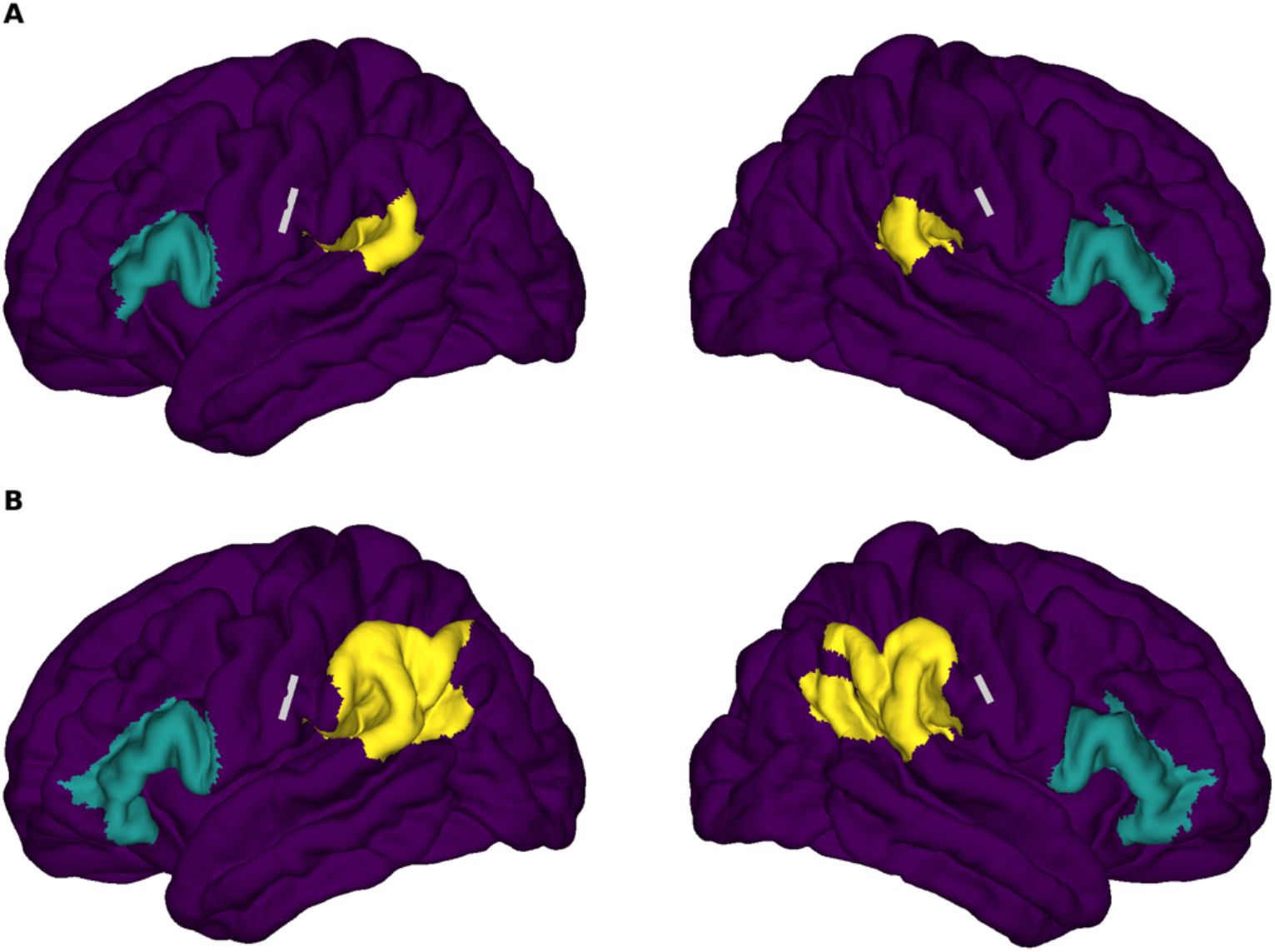
Arcuate / superior lateral fasciculus termination fields. In order to estimate the fraction of tract axons that travel the entire length of the tract, it’s termination fields, centered on Broca’s and Wernicke’s areas, were manually defined in terms of HCP-MMP1.0 parcels according to consensus definitions [1]. (A) Conservative definitions where the anterior termination field, in teal, is composed of parcels 44, 45, 6r, IFSa, IFSp, and FOP4 and the posterior termination field, in yellow, is composed of parcels PSL, RI, STV, and PFcm. For this definition trans-terminal axons account for 0.6% and 0.8% of tract axons in the left and right hemispheres. (B) Liberal definitions in which parcels 47l and p47r were added anteriorly and parcels PF, PFm, and PGi were added posteriorly, resulting in trans-terminal axons accounting for 1.9% and 2.9% of tract axons in the left and right hemispheres. Gray lines indicate the approximate locations at which the tract diameter, used to estimate the total number of tract axons were ascertained [2]. The source data for this figure can be found at https://doi.org/10.5281/zenodo.6097026.

**S5 Fig:**
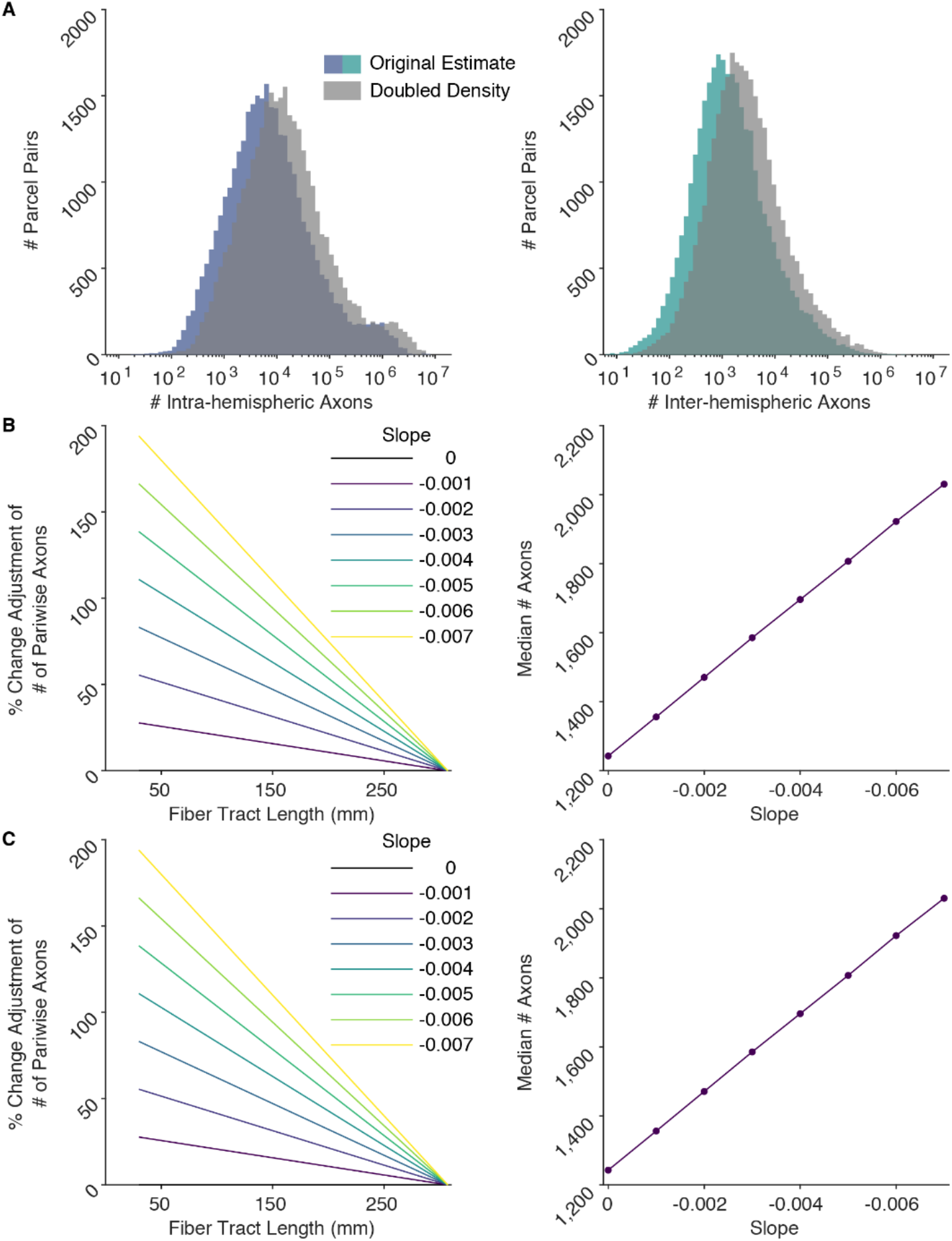
Simulated effect of short axon underestimation on pairwise axon counts. In order to explore the possibility that dMRI tractography is less sensitive to shorter connections the streamline-to-axon ratio was increased linearly with inverse fiber length using a range of slope parameters. Packing density was assumed to be reciprocally decreased in order to fix a constant total cerebral white matter volume. (A) Distributions of Intra- and inter-hemispheric inter-areal axons counts. Gray histograms show the effect of uniformly halfling the ratio and doubling the assumed packing density. (B) intra-hemispheric and (C) inter-hemispheric adjustments to the # of pairwise axons as a function of inverse fiber tract length and the resultant increase in median axon count as a function of the adjustments’ slope parameter. At slope = 0, values are unadjusted from the primary analysis. Large adjustments only increase median counts modestly in the context of the log-normal distribution. For example, for intra-hemispheric connections (B), at a slope of 0.004, the number of the shortest axons is about doubled (i.e., corresponding to doubling the axon density and halving the sensitivity of dMRI to axons), but the median number of inter-parcel axons only increases by ~36%. The source data for this figure can be found at https://doi.org/10.5281/zenodo.6097026.

